# The Development of Sleep/Wake Disruption and Cataplexy as Hypocretin/Orexin Neurons Degenerate in Male vs. Female *Orexin/tTA; TetO-DTA* Mice

**DOI:** 10.1101/2021.10.13.463880

**Authors:** Yu Sun, Ryan Tisdale, Sunmee Park, Shun-Chieh Ma, Jasmine Heu, Meghan Haire, Giancarlo Allocca, Akihiro Yamanaka, Stephen R. Morairty, Thomas S. Kilduff

## Abstract

Narcolepsy Type 1 (NT1), a sleep disorder with similar prevalence in both sexes, is thought to be due to loss of the hypocretin/orexin (Hcrt) neurons. Several transgenic strains have been created to model this disorder and are increasingly being used for preclinical drug development and basic science studies, yet most studies have solely used male mice. We compared the development of narcoleptic symptomatology in male vs. female *orexin-tTA; TetO-DTA* mice, a model in which Hcrt neuron degeneration can be initiated by removal of doxycycline (DOX) from the diet. EEG, EMG, body temperature, gross motor activity and video recordings were conducted for 24-h at baseline and 1, 2, 4 and 6 weeks after DOX removal. Female DTA mice exhibited cataplexy, the pathognomonic symptom of NT1, by Week 1 in the DOX(-) condition but cataplexy was not consistently present in males until Week 2. By Week 2, both sexes showed an impaired ability to sustain long wake bouts during the active period, the murine equivalent of excessive daytime sleepiness in NT1. Body temperature appeared to be regulated at lower levels in both sexes as the Hcrt neurons degenerated. During degeneration, both sexes also exhibited the “Delta State”, characterized by sudden cessation of activity, high delta activity in the EEG, maintenance of muscle tone and posture, and the absence of phasic EMG activity. Since the phenotypes of the two sexes were indistinguishable by Week 6, we conclude that both sexes can be safely combined in future studies to reduce cost and animal use.

**Statement of Significance:** Although narcolepsy is a disorder that affects both men and women with similar frequency, most basic research and preclinical development studies of sleep have utilized male experimental subjects. The identification of the hypocretin/orexin (Hcrt) neuron loss as the likely cause of human narcolepsy has led to the development of transgenic mouse strains that model this disorder. Here, we compare the emergence of narcoleptic symptoms in male vs. female bigenic *orexin-tTA; TetO DTA* mice, a state-of-the-art narcolepsy model in which degeneration of the Hcrt neurons can be triggered by dietary manipulation. We find that female mice develop the narcoleptic phenotype more rapidly than males but that both sexes are equally symptomatic by the end of the degeneration period.

## Introduction

Narcolepsy, a lifelong illness characterized by excessive daytime sleepiness (EDS) and associated symptoms, is classified as type 1 or 2 (NT1 and NT2) based on the presence or absence of cataplexy and/or hypocretin/orexin deficiency, with up to 60% of patients with narcolepsy having NT1^1-3^. Approximately 1 in 2000 people are diagnosed with narcolepsy^4-7^ and more than 50% of patients report that their first symptoms occurred before 16 years of age^2,8^. However, only 18% of patients with narcolepsy receive a diagnosis within 1 year of symptom onset and only 50% of patients with narcolepsy receive a diagnosis within 5 years^2,9^.

Conflicting reports exist regarding sex differences in the prevalence of narcolepsy. Epidemiological studies have indicated that narcolepsy is more common in men than in women^4,6,10^. When the research focus is restricted to NT1, however, either no sex difference is reported^5,11-13^ or the incidence is reported as higher in women^6^. Due to overall differences in access to health services, women are less likely to undergo polysomnographic assessment^14^, so the existing literature may not be completely reliable. Nonetheless, cataplexy has been reported to be more common in women than in men^5,15^, although symptom severity may not differ between the sexes ^13^.

Since NT1 is a debilitating disorder, pharmacological management of symptoms is critical to maintain quality of life.^16^ With the development of mouse models of narcolepsy that better reflect the loss of hypocretin/orexin (Hcrt) neurons that underlie this disorder^17^, future drug development for narcolepsy will likely rely on such animal models^18^. Indeed, there has been increasing use on such models for both basic science and preclinical development studies in recent years^19-26^. In such studies, however, only male mice have typically been used, which is problematic for several reasons. First, use of a single sex ignores biology that is relevant to the at least half the human population, particularly since baseline differences in sleep have been documented in both humans and rodents^27^. Furthermore, a precedent already exists for a difference in recommended doses for a sleep medication in men vs. women^28^. Lastly, breeding of transgenic mice is expensive and time-consuming and to discard half of the offspring violates one of the 3R principles^29^.

In the present study, we compare the development of narcolepsy symptoms in male and female bigenic *orexin-tTA; TetO-DTA* mice, a state-of-the-art narcolepsy model in which degeneration of the Hcrt neurons can be initiated by a simple dietary manipulation^30^. We find that female mice progress into the narcoleptic phenotype more rapidly than male mice but that both sexes are equally symptomatic by the end of the degeneration period. We also find a novel state that emerges in both sexes during Hcrt neuron degeneration that we call the “Delta State”, which is characterized by sudden, brief periods of movement cessation without a loss of muscle tone that is accompanied by high delta activity in the EEG while the eyes remain open.

## Methods

All experimental procedures were approved by the Institutional Animal Care and Use Committee at SRI International and were conducted in accordance with the principles set forth in the *Guide for Care and Use of Laboratory Animals*.

### Animals

“DTA mice” (N=7 males; N=7 females) were the double transgenic offspring of *orexin/tTA* mice (C57BL/6-Tg(*orexin/tTA*)/Yamanaka)^30^, which express the tetracycline transactivator (tTA) exclusively in Hcrt neurons^31^, and B6.Cg-Tg(*tetO-DTA*)1Gfi/J mice^32^ (JAX #008468), which express a diphtheria toxin A (DTA) fragment in the absence of dietary doxycycline (DOX). Both parental strains were from a C57BL/6J genetic background. Parental strains and offspring used for EEG/EMG recording were maintained on a diet (Envigo T-7012, 200 DOXycycline) containing doxycycline (DOX(+) condition) to repress transgene expression until neurodegeneration was desired. Hcrt neuron degeneration was initiated by substituting DOX(+) chow with normal rodent chow (DOX(-) condition). When the diet was changed from DOX(+) to DOX(-) condition, the animal’s cage was changed as well to prevent continued DOX exposure due to coprophagy. Mice were maintained on normal chow for 6 weeks and then DOX(+) chow was reintroduced to arrest any further neurodegeneration. All mice were maintained on a LD12:12 light:dark cycle at room temperature (22±2°C; 50±20% relative humidity), had access to food and water *ad libitum*, were weighed weekly.

### Surgical Procedures

Because of the differential growth rate of males and females (**Figure 1**), the two sexes of DTA mice underwent surgery at slightly different ages. To ensure that all mice were at least 20 g at the time of surgery (the minimum recommended body mass for this procedure), male mice were implanted at 9±1 weeks of age and were 25.7±0.5 g, whereas female DTA mice were implanted at 14±1 weeks and were 20.7±0.8 g. Mice were anesthetized with isoflurane and sterile telemetry transmitters (HD-X02 for males and F20-EET for females, Data Sciences Inc., St Paul, MN) were placed subcutaneously. Biopotential leads were routed subcutaneously to the head and EMG leads were positioned in the right nuchal muscle. Cranial holes were drilled through the skull at -2.0 mm AP from bregma and 2.0 mm ML and on the midline at -1 mm AP from lambda. The two biopotential leads used as EEG electrodes were inserted into these holes and affixed to the skull with dental acrylic. The incision was closed with absorbable suture. Analgesia was managed with meloxicam (5 mg/kg, s.c.) and buprenorphine (0.05 mg/kg, s.c.) upon emergence from anesthesia and for the first day post-surgery. Meloxicam (5 mg/kg, s.c., q.d.) was continued for 2 d post-surgery.

**Figure 1.**
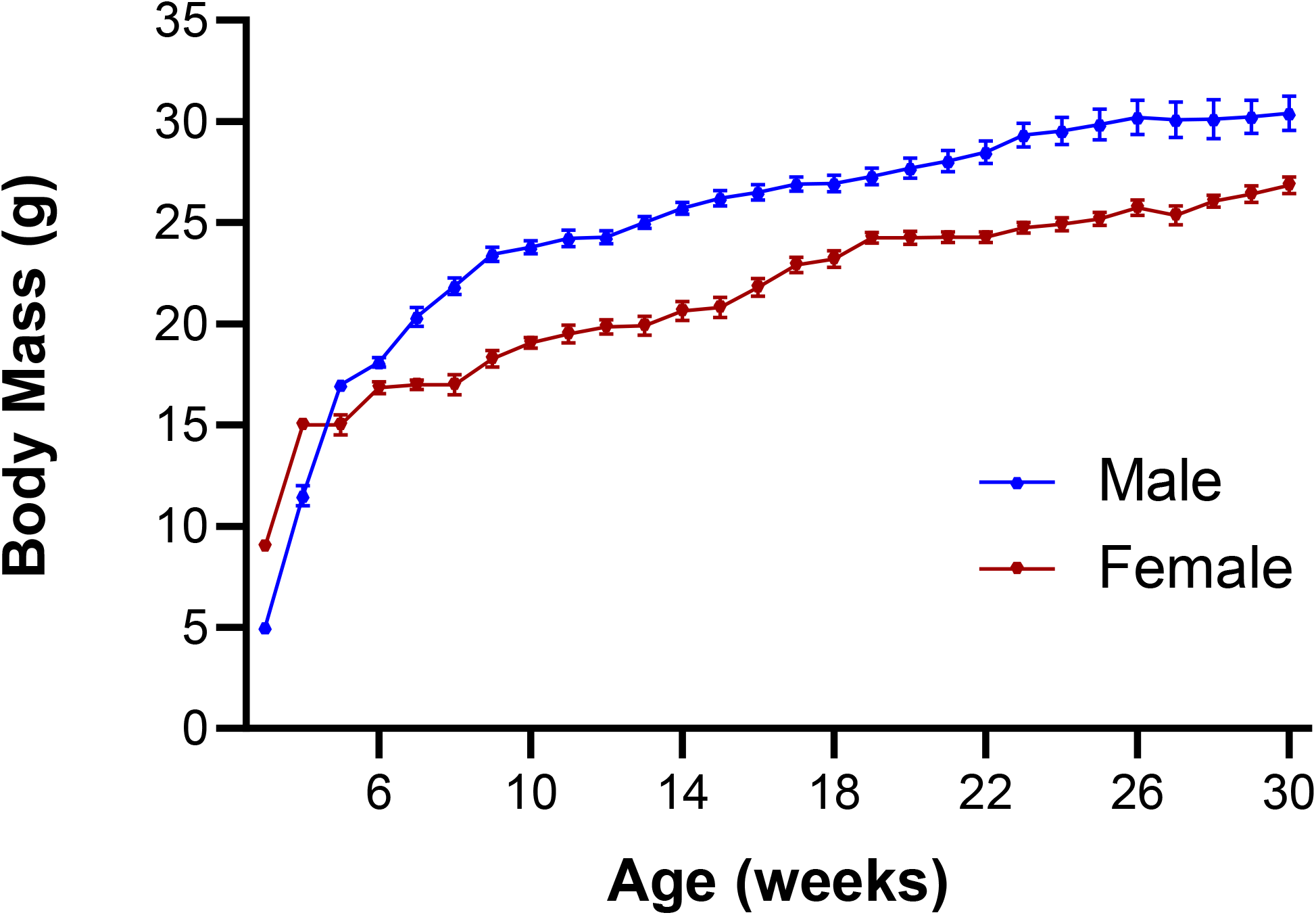
Growth curves for male (N = 18) and female (N = 14) *orexin/tTA; TetO-DTA* mice. Values are mean ± SEM.

### EEG, EMG, activity and body temperature recording

Prior to data collection, DTA mice had 2 weeks post-surgical recovery and at least 2 weeks adaptation to running wheels. All mice then underwent a 24 h baseline (Week 0) recording in which digital videos, EEG, EMG, subcutaneous body temperature (T_b_), and gross motor activity were recorded via telemetry using Ponemah (DSI, St Paul, MN). Digital videos were recorded at 10 frames per second, 4CIF de-interlacing resolution; EEG and EMG were sampled at 500 Hz. Mice were then switched to normal chow (DOX(-) condition) to induce expression of the DTA transgene specifically in the Hcrt neurons and thereby initiate degeneration of these cells^30^ and 24 h recordings were again conducted at 1, 2, 4 and 6 weeks in the DOX(-) condition. To ensure that data collection occurred at the same phase of the estrus cycle for females, 24 h recordings occurred at 4 day intervals, e.g., on day 8 in the DOX(-) condition during Week 1, on day 16 DOX(-) during Week 2, on day 28 during Week 4, and on day 42 during Week 6. After completion of the recording at 6 wk, DOX chow was reintroduced to minimize further degeneration. Tail snips to confirm the DTA genotype were obtained at weaning and at the end of the study.

Subsequent to surgery, mice were housed individually in home cages with access to food, water, nestlets and running wheels *ad libitum*. Room temperature, humidity, and lighting conditions (LD12:12; lights on at 06:00) were monitored continuously. Animals were inspected daily in accordance with AAALAC and SRI guidelines and body weights were taken weekly.

### Classification of Arousal States

For all recordings, video, EEG and EMG data collected during the first 6-h of the dark period (ZT13-ZT18) were used by expert scorers to classify 10-s recording epochs as wakefulness (W), non-Rapid Eye Movement (NREM) sleep, REM sleep, or cataplexy using NeuroScore (DSI, St. Paul, MN) as in our previous studies^19,20^. Criteria for cataplexy were ≥ 10 s of EMG atonia, theta-dominated EEG, and video-confirmed behavioral immobility preceded by ≥40 s of wakefulness^33^. Running wheel activity, determined from video recordings, was scored in 10-s epochs for the purpose of training Somnivore (see below). While analyzing these recordings, we recognized a novel state that we called the Delta State (DS). Criteria for scoring an epoch as DS were the sudden cessation of locomotion during W, the persistence of muscle tone but the absence of phasic EMG activity, and a synchronous, NREM-like EEG pattern while the eyes remained open and the mouse remained in a standing position. DS terminated abruptly and mice rapidly resumed ambulatory behaviors with typical Wake EEG patterns.

### Application of a Supervised Machine Learning Model to Assist in Arousal State Classification

To score the remaining 18-h of each 24-h recording, the manually scored 6-h period of each recording (ZT13-ZT18) was provided to Somnivore (ver. 1.0.70)^34^ in EDF format to create a training dataset and for subsequent assessment of autoscoring accuracy (see below). From the manually scored 6-h recordings of each mouse, 100 10-s epochs of Wake and 100 epochs of running wheel activity were randomly selected to train a classifier. If a recording had fewer than 100 epochs of running wheel activity in the initial 6-h period, additional epochs were selected from the remaining 18-h recording. Since little NREM and REM occurred during the first 6-h of the active phase (ZT13-ZT18), training epochs for these states were manually selected from throughout the entire 24-h recording. For cataplexy, if the 6-h recordings contained fewer than 100 epochs, REM and cataplexy were trained as a single state and were manually separated post-autoscoring through application of the scoring rules described below. The selected training epochs for all states were then applied in a supervised machine learning model to automatically score sleep-wake states for the entire 24-h period.

Following initial autoscoring, agreement with the manually scored 6-h period was assessed and generalization of the autoscoring to the remaining 18-h of the recording was evaluated. An additional 5-25 epochs of states with low accuracy values were added to the training dataset and recordings were automatically scored again. Following this optimization phase, scoring rules were applied to the Somnivore-scored dataset to select potentially misidentified epochs of cataplexy and/or REM. Manual correction of the scoring occurred when autoscoring provided any of the following unlikely results:

- single epochs of REM
- epochs of REM preceded by W, cataplexy or wheel running
- epochs of cataplexy preceded by NREM or REM
- prolonged or frequently occurring short bouts of cataplexy or REM

Finally, accuracy of autoscoring was evaluated by comparing the corrected autoscoring to the manual 6-h scoring with a function provided by the software that enables comparison of the *F* value for each state, a measure of precision and accuracy used to quantify algorithm generalization^34^. Accuracy of the autoscored data had high concordance with manual scoring, with *F* values for most states indicating a high level of accuracy.

### Data analysis and statistics

Among the 7 female DTA mice implanted for this study, the quality of the recordings was problematic for one mouse on Week 0 and another female on Week 6. Due to the repeated measures experimental design of this study, it was necessary to eliminate these two females from the data analysis; thus, statistical analyses are based on N=7 males and N=5 females. In the Results and Discussion below, however, these two mice are included for descriptive rather than quantitative statements regarding the presence or absence of cataplexy during Weeks 1-4.

For all states, data were analyzed as time spent in state per hour and cumulative time spent in each state. Sleep/wake architecture measures included the duration and the number of bouts for each state. A “bout” of a particular state was defined as 2 or more consecutive epochs of that state and ended with a single epoch of any other state.

The EEG power spectrum (0.5-100 Hz) during W, NREM and REM was analyzed by fast Fourier transform algorithm on all artifact-free epochs. For spectral analyses, a minimum of 6 consecutive epochs of Wake or NREM and a minimum of 3 consecutive epochs of REM was required for inclusion in the analysis. For cataplexy and Delta State, there was no minimum number of epochs criterion; all epochs were included in spectral analyses. For each mouse, EEG power for W, NREM and REM during the degeneration period was normalized to the 6-h average power per bin during the pre-degeneration baseline recording (Week 0). To determine whether any changes in EEG power occurred across the degeneration phase, we compared EEG power within each state across DOX(-) weeks to the DOX(+) condition. Since cataplexy did not occur during the baseline recording in the DOX(+) condition, Wake EEG power during Week 0 was used for normalization of cataplexy across degeneration weeks. EEG spectra for Wake and Cataplexy were analyzed in 0.122 Hz bins and in standard frequency bands rounded to the nearest half-Hz value (delta: 0.5-4Hz, theta: 6-9Hz, alpha: 9-12Hz, beta: 12-30Hz, low gamma: 30-60Hz and high gamma: 60-100 Hz). Hourly averages of T_b_ and gross activity determined from the telemetry transmitters were also analyzed.

Total time in states, REM/NREM ratios, average bout durations, average number of bouts, and standard frequency EEG power bands were analyzed using 1-way repeated-measures analysis of variance (ANOVA). Due to continuous wakefulness during the 12-h dark period during Week 0 in one female DTA mouse and the resultant absence of NREM, REM, cataplexy and related EEG power bands for that individual, mixed-effect analysis was used in some analyses to compensate for the missing data. When 1-way ANOVA or mixed-effect analysis indicated statistical significance, the *post hoc* Dunnett multiple comparisons test was performed to determine specific differences. Because the continuous wakefulness during the dark period during Week 0 in the female mouse mentioned above resulted in a single long Wake bout duration, the Friedman test was applied to analyze female Wake bout duration. All other data analyses were by 2-way repeated-measures ANOVA within each sex with treatment (DOX(-) week) and time of day as factors. When 2-way ANOVA indicated statistical significance, paired two-tailed *t*-tests were performed *post hoc* to determine specific differences. Statistics were calculated using the functions provided in the MATLAB statistics and machine learning toolbox and using GraphPad Prism (ver. 8.4.2). For the EEG spectra data, statistical comparisons were calculated only on the standard frequency bands.

## Results

### State Amounts and the Development of Cataplexy

Relatively few changes were observed for hourly, total and cumulative time spent in Wake, NREM or REM during the 24 h L/D cycle across the 6 week degeneration period (**Figures 2, 3**). Wake decreased in male DTA mice across the 24 h period during Week 6 of the degeneration (**Figure 2A**); this decrease was primarily due to a reduction in Wake during the first half of the dark phase. Total time in Wake decreased in the dark phase during Week 6 but increased slightly in the light phase (**Figures 2C, 2D**). Cumulative time in Wake decreased for both Week 4 and Week 6 in males (**Figure 3A**). In females, Wake decreased across the 24 h cycle during Week 4 due primarily to a decrease in the dark phase that was also evident in Week 6 (**Figures 2B, 2C**). Cumulative time in Wake also decreased for both Week 4 and Week 6 in females (**Figure 3B**).

**Figure 2.**
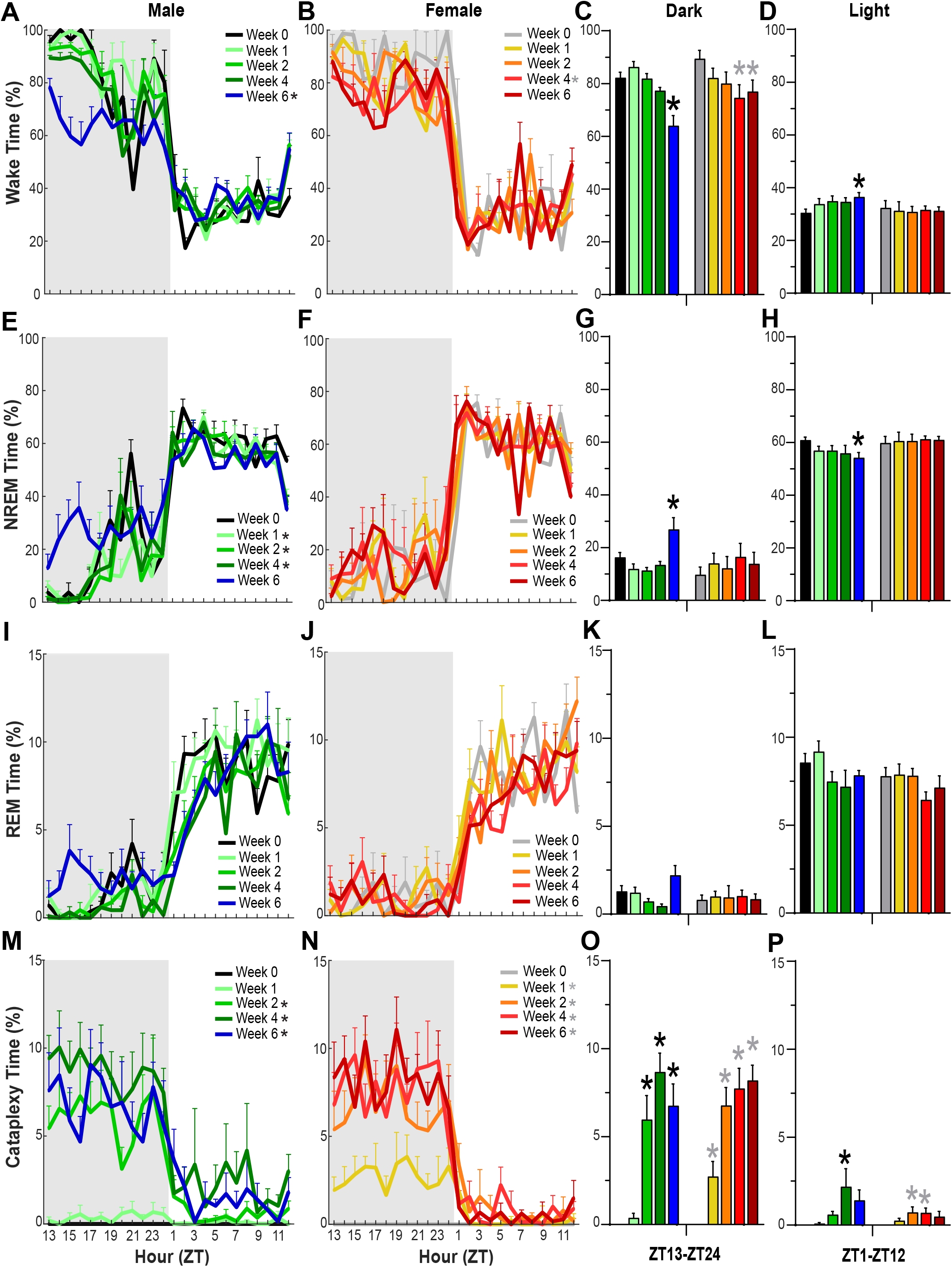
Hourly percentage of time in Wakefulness, NREM sleep, REM sleep, and cataplexy in narcoleptic male (**A, E, I, M**) and female (**B, F, J, N**), *orexin/tTA; TetO-DTA* mice during baseline (0 Week) and Weeks 1, 2, 4 and 6 in the DOX(-) condition during which time the Hcrt neurons are expected to degenerate. * in the legend within each panel indicates a significant difference (*p* < 0.05) during that week relative to baseline Week 0 as determined by ANOVA. Panels **C, G, K** and **O** summarize the percentage of time that each sex spent in these states during the 12-h dark period whereas panels **D, H, L** and **P** summarize these data for both sexes during the 12-h light period. * above the bars in each panel indicate significance (*p* < 0.05) during that week relative to baseline Week 0 as determined by ANOVA. Values are mean ± SEM.

**Figure 3.**
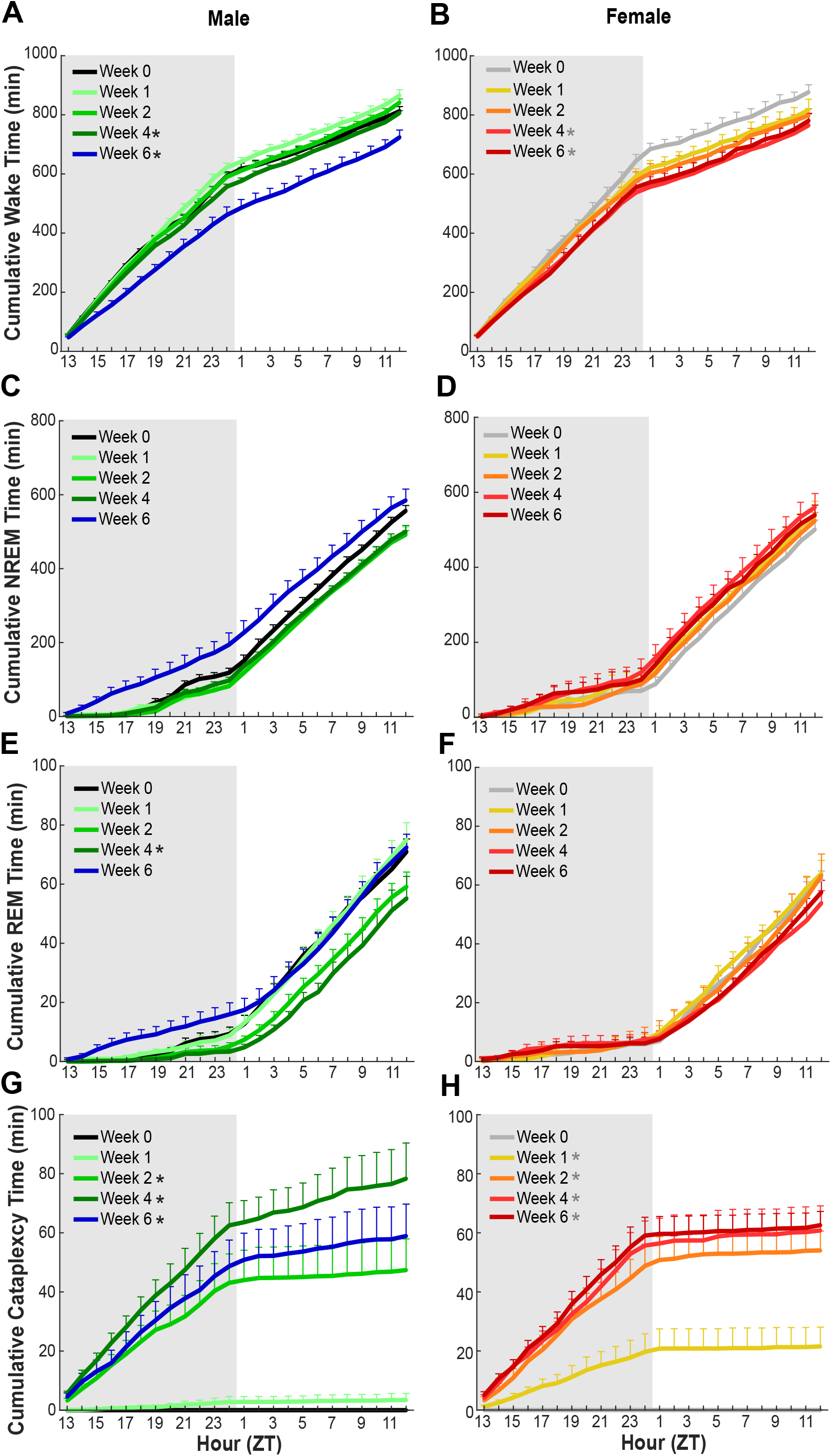
Cumulative amounts of Wakefulness, NREM sleep, REM sleep, and Cataplexy in narcoleptic male **(A, C, E, G)** and female **(B, D, F, H)** *orexin/tTA; TetO-DTA* mice during baseline (0 Week) and Weeks 1, 2, 4 and 6 in the DOX(-) condition during which time the Hcrt neurons are expected to degenerate. Shaded area indicates the dark period of the 24-h cycle. Values are mean ± SEM. * in the legend indicates a significant difference (*p* < 0.05) during that week relative to baseline Week 0 as determined by ANOVA.

NREM decreased overall in males during Weeks 1-4 and in the light phase during Week 6 (**Figures 2E and 2H**). In contrast, NREM increased in the dark phase during Week 6 (**Figure 2G**). No significant differences were found for NREM in females across the 6 week degeneration period (**Figures 2F, 2G and 2H**). For REM, no significant differences were observed in either males or females for percent time or total time across the 24-h period (**Figures 2I-2L**). However, cumulative time decreased overall for Week 4 in male mice (**Figure 3E**).

The emergence of cataplexy during the degeneration period differed between male and female DTA mice (**Figures 2M-2P, 3G and 3H**). Cataplexy was evident in all 7 female mice during the first week of the degeneration while only 3 of 7 males had discernible bouts. However, by Week 2 and for subsequent weeks, all males and females exhibited cataplexy. Cataplexy significantly increased across the 24 h recordings and cumulatively for Weeks 2-6 in males and for Weeks 1-6 in females. For both sexes, cataplexy was most evident in the dark (active) phase. In males, total cataplexy time increased during the dark phase in Weeks 2-6 and during the light phase in Week 4 whereas, in females, total cataplexy time increased during the dark phase in Weeks 1-6 and during the light phase in Weeks 2-4 (**Figure 2O, 2P**).

Neither the NREM nor REM sleep latencies (measured from lights off) nor REM:NR ratios were significantly different from Week 0 for any degeneration week (data not shown).

### Sleep/Wake Architecture Measures

As indicated by both the progressive reduction in mean Wake bout duration (**Figures 4A, 4B)** and the increased number of Wake bouts (**Figures 5A, 5B)**, Wakefulness became more fragmented in both female and male DTA mice during Hcrt neuron degeneration but the phenotype differed somewhat. In the dark phase, mean Wake bout duration decreased during Weeks 2-6 in both sexes whlie the number of Wake bouts increased during Weeks 2-6 in males and Weeks 1-6 in females. However, in the light phase, mean Wake bout duration decreased and the number of Wake bouts increased in male mice for Weeks 1-6 whereas, for female mice, the number of Wake bouts increased during Week 1 without any significant change in bout duration.

**Figure 4.**
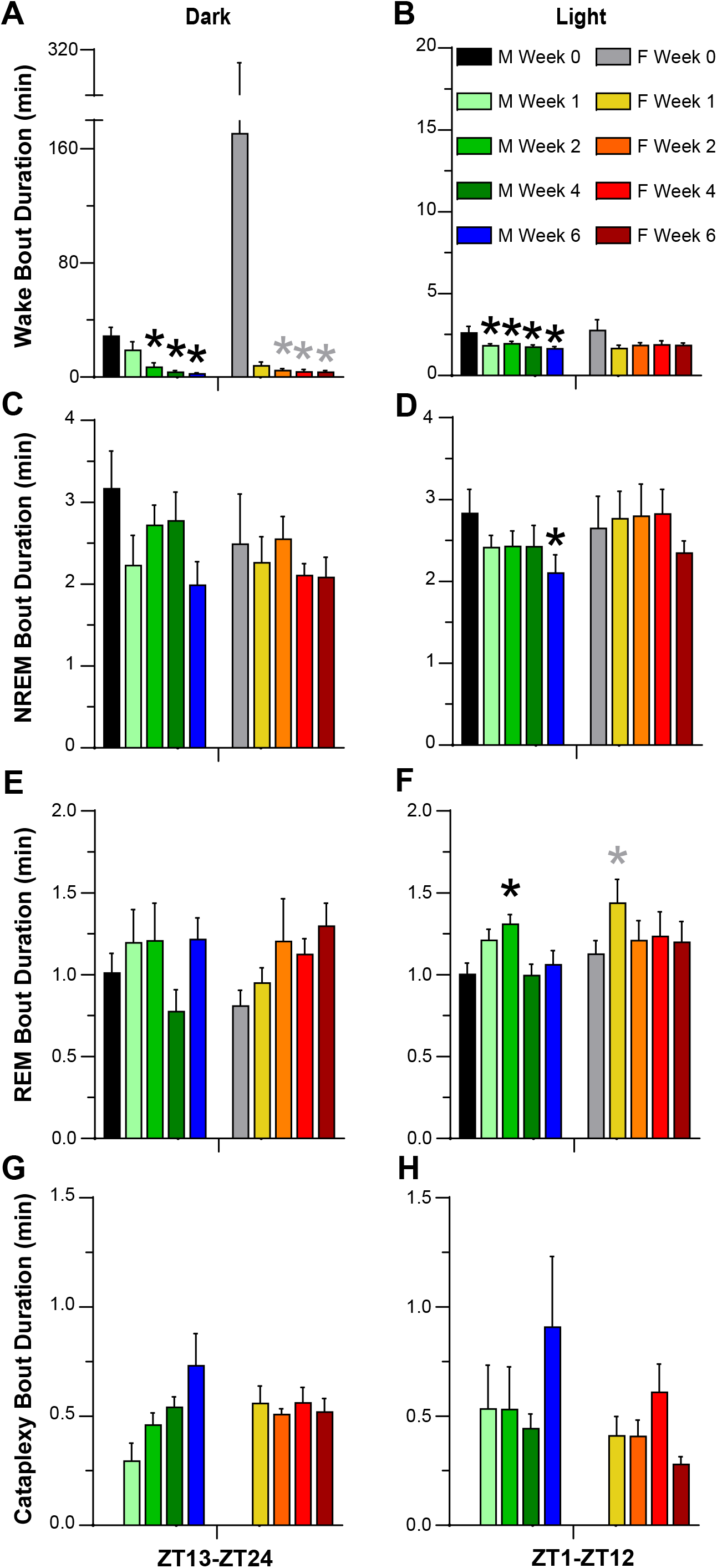
Mean bout durations for Wakefulness (**A, B**), NREM sleep (**C, D**), REM sleep (**E, F**), and cataplexy (**G, H**), during the 12-h dark (left) and light (right) periods for narcoleptic male and female *orexin/tTA; TetO-DTA* mice during baseline (Week 0) and Weeks 1, 2, 4 and 6 in the DOX(-) condition during which time the Hcrt neurons are expected to degenerate. Values are mean +/-SEM. * above the bars in each panel indicate significance (*p* < 0.05) during that week relative to baseline Week 0 as determined by ANOVA or the Friedman test, which was applied to analyze Wake bout duration in female mice due to the one female who did not sleep at all during the dark period. Values are mean ± SEM.

**Figure 5.**
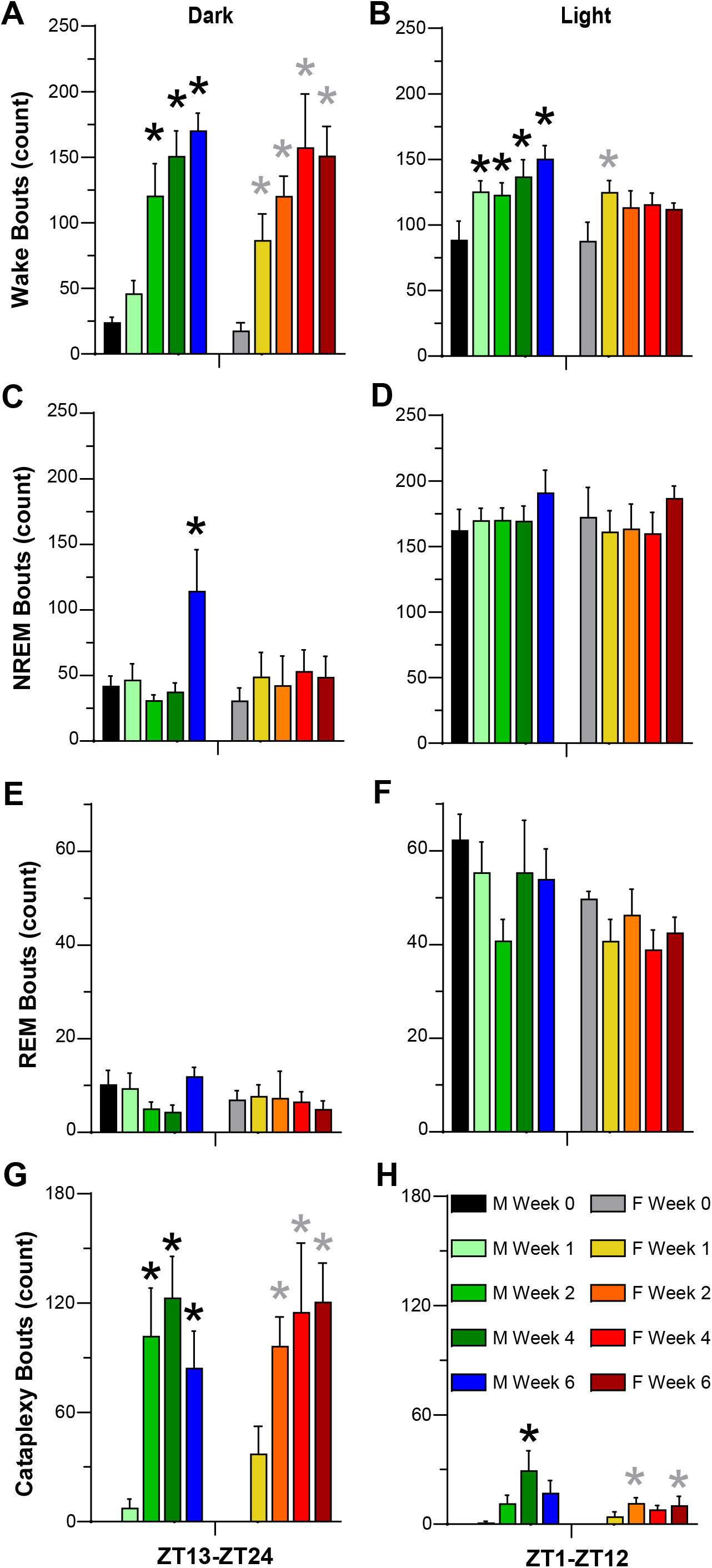
Number of bouts of Wakefulness (**A, B**), NREM sleep (**C, D**), REM sleep (**E, F**), and cataplexy (**G, H**), during the 12-h dark (left) and light (right) periods for narcoleptic male and female *orexin/tTA; TetO-DTA* mice during baseline (Week 0) and Weeks 1, 2, 4 and 6 in the DOX(-) condition during which time the Hcrt neurons are expected to degenerate. Values are mean ± SEM. * above the bars in each panel indicate significance (*p* < 0.05) during that week relative to baseline Week 0 as determined by ANOVA. Values are mean ± SEM.

NREM bout duration decreased in the light phase (**Figure 4D**) and the number of NREM bouts increased during the dark phase (**Figure 5C**) during Week 6 for male DTA mice; no change in these parameters were observed in females. REM bout duration increased in the light phase during Week 2 in male mice and during Week 1 in female mice (**Figure 4F**). Cataplexy bout duration during the dark phase appeared to increase progressively with Hcrt neuron degeneration in male but not female DTA mice (**Figure 4G**). The number of cataplexy bouts increased in the dark phase for both sexes during weeks 2-6 and during the light phase in Week 4 for males and Weeks 2 and 6 for females (**Figure 5G, 5H**).

### Activity and Body Temperature Rhythms

Gross motor activity increased in males for Weeks 2-4 but were at baseline levels on Week 6 (**Figure 6A**). No significant differences in gross activity were observed in females (**Figure 6B**), perhaps due to the high variation during Week 0. T_b_, measured subcutaneously, decreased during Weeks 2-6 in both male and female mice (**Figures 6C, 6D**). This decrease occurred across the 24 h period and was indicative of an overall reduction of subcutaneous T_b_ during both the dark and light phases. For males, mean T_b_ declined significantly in both the dark and light phases during Weeks 2-6 (**Figure 6E**). For females, significant reductions in T_b_ occurred for Week 4 during the dark and for Weeks 2-6 during the light phase (**Figure 6F**).

**Figure 6.**
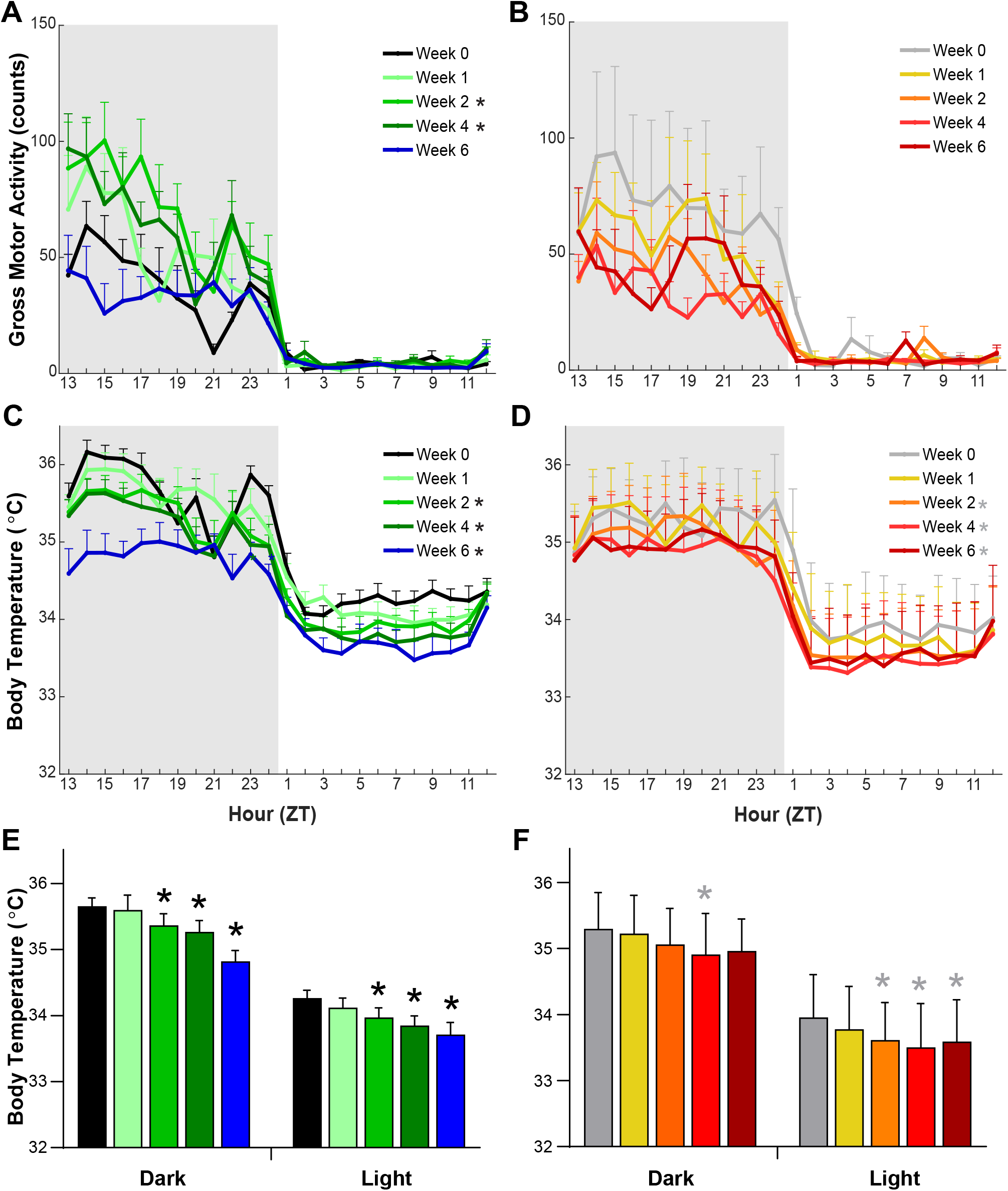
Gross motor activity (**A, B**) and subcutaneous body temperature (**C, D**) in narcoleptic male (left) and female (right) *orexin/tTA; TetO-DTA* mice during baseline (0 Week) and weeks 1, 2, 4 and 6 in the DOX(-) condition during which time the Hcrt neurons are expected to degenerate. Mean subcutaneous body temperature during the 12-h dark and 12-h light for males (**E**) and females (**F**). Values are mean ± SEM. * in the legend and above the bars indicates a significant difference (*p* < 0.05) during that week relative to baseline Week 0 as determined by ANOVA.

### EEG Power Spectra

Spectral analyses were only performed on the manually-scored first 6-h of the dark phase for each recording since this period was carefully evaluated for artifact removal. Analyses during this period revealed progressive changes in Wake EEG which was more evident in males than females (**Figure 7**). When compared to pre-degeneration Wake (normalized to Week 0), increased spectral power was observed from 2-9 Hz and 15-25 Hz for both sexes (**Figures 7A, 7B**). However, when EEG power was binned into the standard power bands, few statistical differences were found (**Figures 7C-7H**). Delta power increased in males during Week 4 and in females during Weeks 2 and 4, theta power increased in males during Week 4, and beta power increased in males during Weeks 2-4 and in females during Weeks 1-6. Since the F20-EET transmitters used in the female mice have an attenuated signal above 60 Hz, high gamma results for female mice (**Figure 7H**) are included for informational purposes only although the statistical results obtained are consistent with the decline in high gamma observed in males during Week 6.

**Figure 7.**
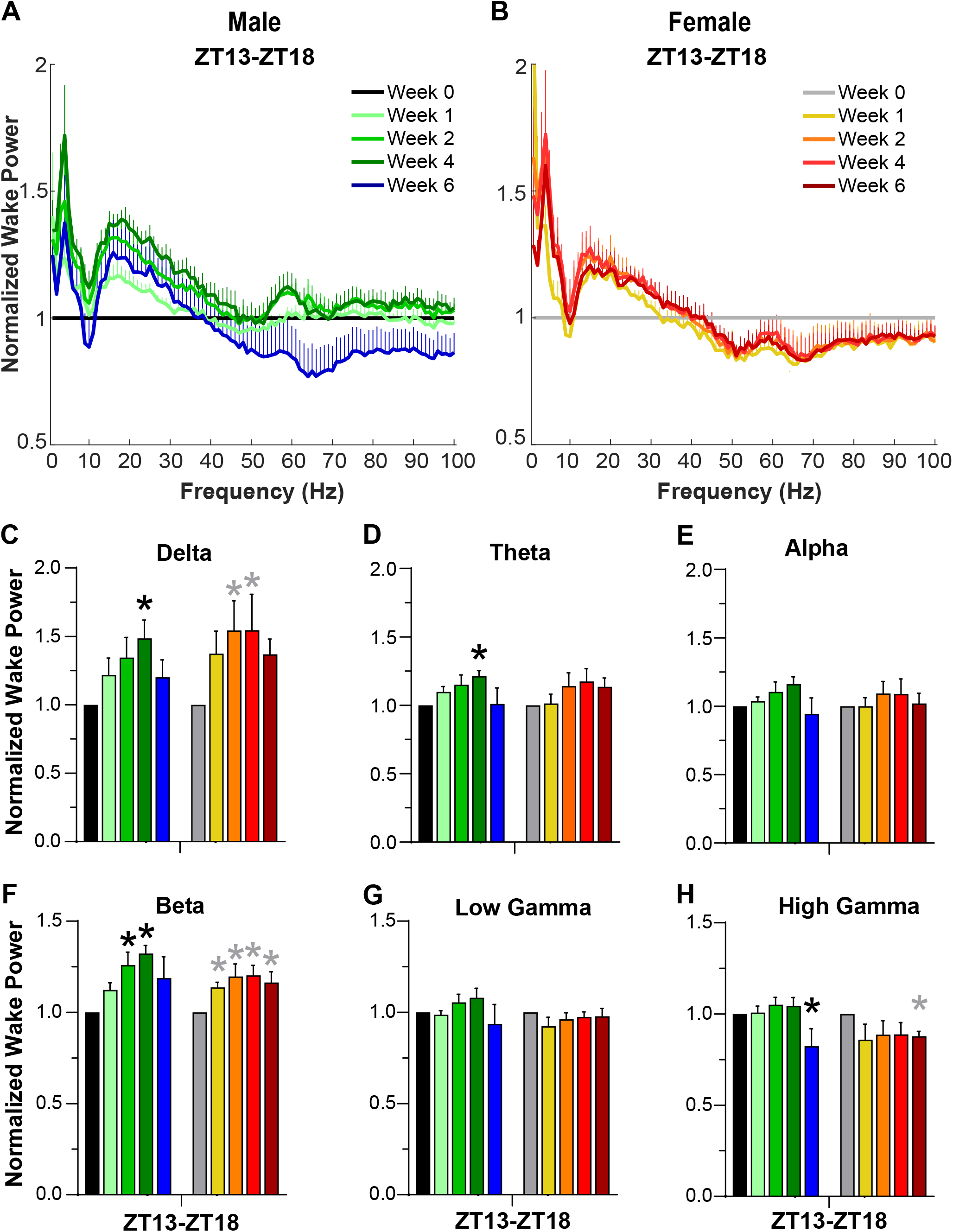
Normalized EEG spectral power (0-100 Hz) during wakefulness for (**A**) male and (**B**) female *orexin/tTA; TetO-DTA* mice during baseline (Week 0) and Weeks 1, 2, 4 and 6 in the DOX(-) condition during which time the Hcrt neurons are expected to degenerate. Normalized Wake Power in male and female *orexin/tTA; TetO-DTA* mice in the (**C**) delta, (**D**) theta, (**E**) alpha, (**F**) beta, (**G**) low gamma and (**H**) high gamma bandwidths. Values are mean ± SEM. * above the bars in panels C-H indicate significance (*p* < 0.05) during that week relative to baseline Week 0.

To evaluate EEG power during cataplexy, we normalized to the average Wake power during Week 0 (**Figure 8**). Increased power in the Delta, theta, alpha and beta ranges were observed with small changes in low and high gamma. Statistical analyses were not performed since no Cataplexy occurred during Week 0.

**Figure 8.**
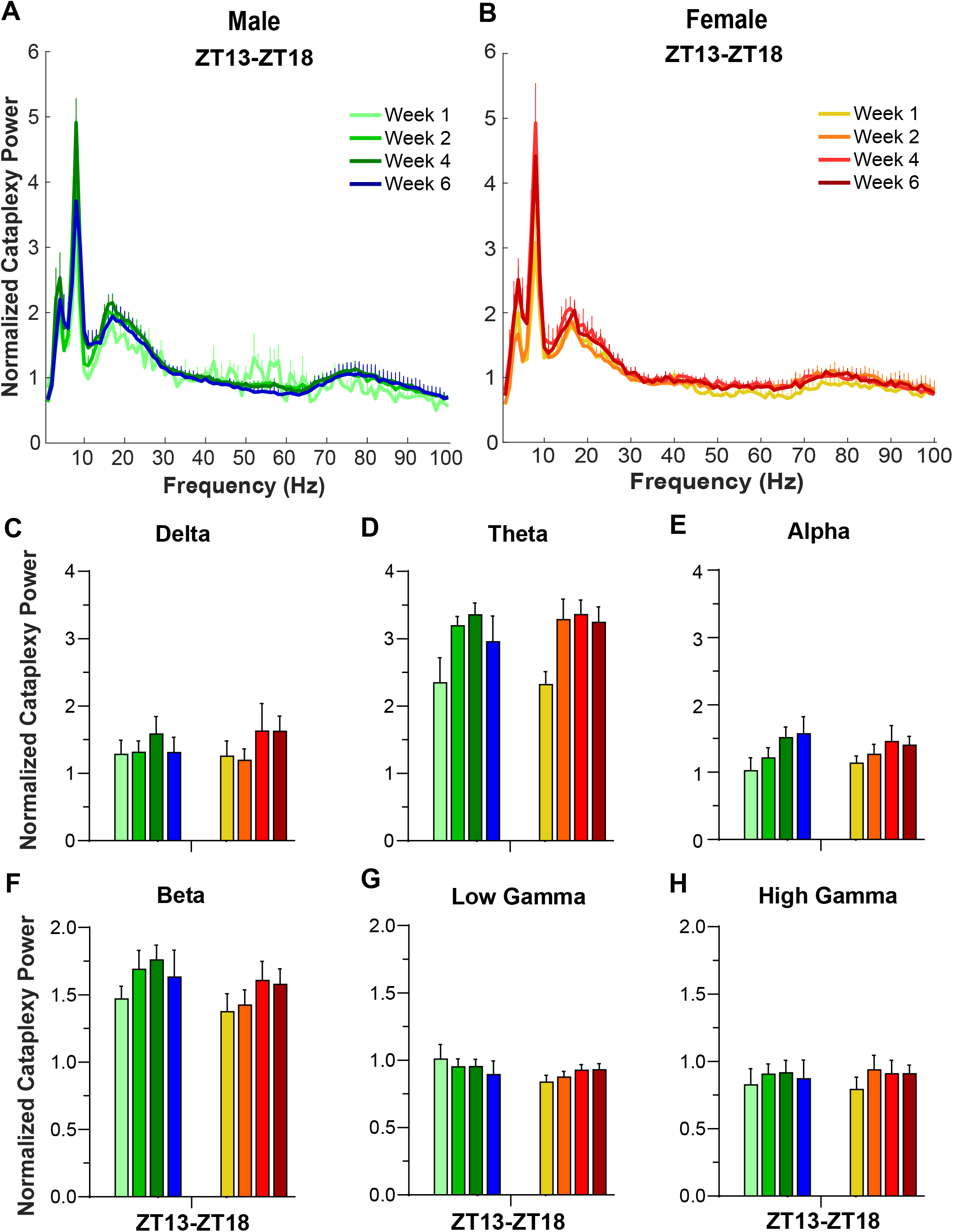
Normalized EEG spectral power (0-100 Hz) during cataplexy for (**A**) male and (**B**) female *orexin/tTA; TetO-DTA* mice during baseline (Week 0) and Weeks 1, 2, 4 and 6 in the DOX(-) condition during which time the Hcrt neurons are expected to degenerate. Normalized EEG power during cataplexy in male and female *orexin/tTA; TetO-DTA* mice in the (**C**) delta, (**D**) theta, (**E**) alpha, (**F**) beta, (**G**) low gamma and (**H**) high gamma bandwidths.

EEG power in NREM and REM was not evaluated due to the paucity of these states during the ZT13-18 time period for most of the mice studied.

### A Novel High EEG Delta State During Wakefulness

Manual scoring of the EEG records revealed a novel state which we labelled as Delta State (DS; **Figure 9**). DS only occurred following the initiation of the degeneration process and became more prevalent as the degeneration proceeded. DS was observed during Wake, typically during active Wake, and was characterized by a sudden cessation of movement with an absence of phasic EMG activity. Differing from cataplexy, muscle tone was not lost and video recordings revealed postural maintenance. Concurrently, the EEG becomes synchronized with high delta activity as in NREM sleep. Without video recordings, these episodes would be difficult to distinguish from NREM. However, the behavior of the mice was distinctly different from that of a mouse entering sleep. Typical behavior that precedes sleep, such as lying down or curling up into their nest, did not occur. The mice were typically ambulatory but suddenly stopped, the EEG became synchronized with high Delta, and no phasic EMG activity occurred. Videos revealed that the eyes remained open. Within 1-6 epochs, the mouse suddenly returned to its ambulatory behavior, a Wake-like EEG reappeared and phasic EMG activity was evident (**Figures 9A-D, Supplemental videos S1 & S2**).

**Figure 9.**
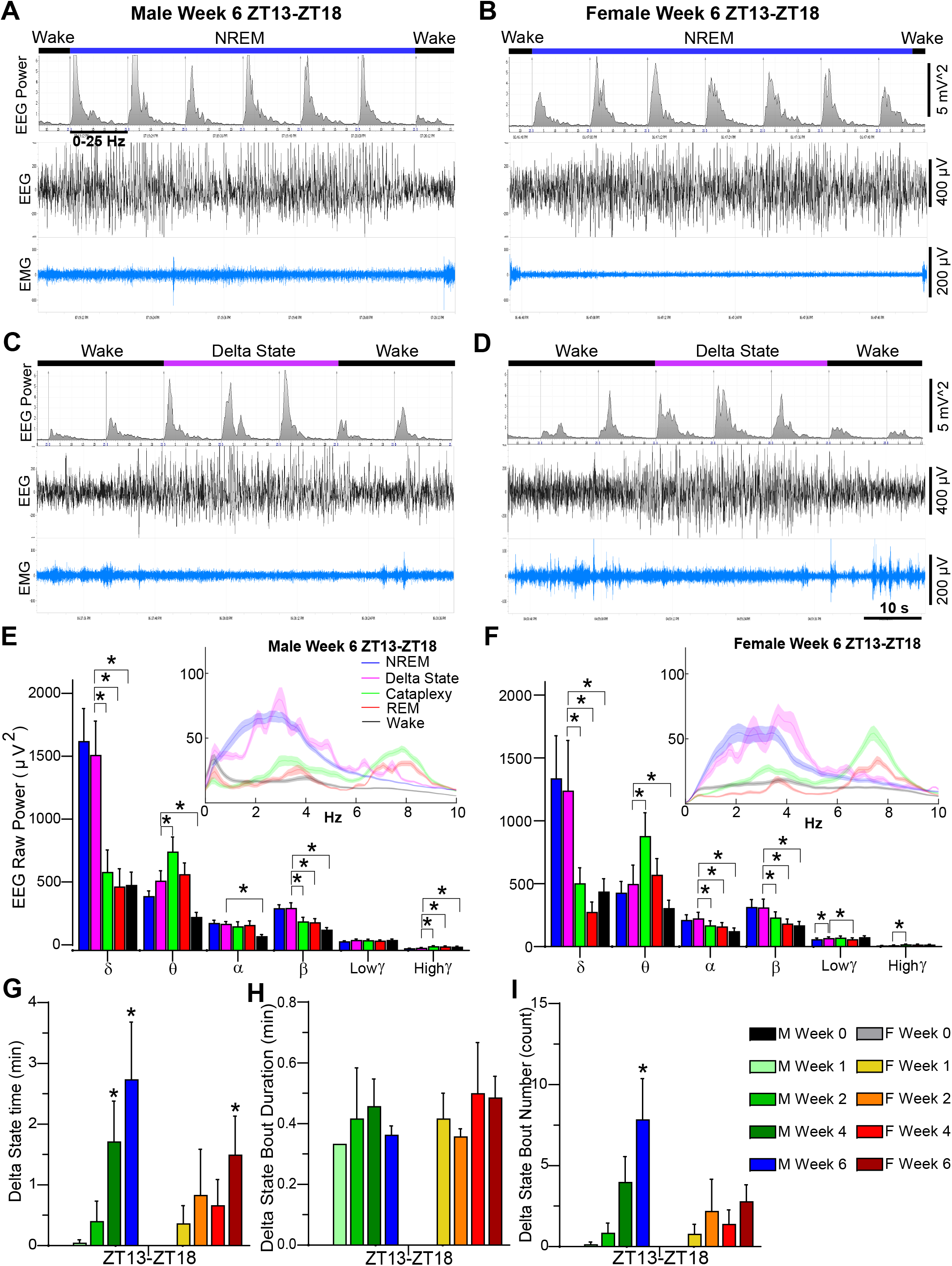
Example EEG spectrograms (7 10-sec epochs), EEG and EMG traces illustrating typical transitions from Wake to NREM sleep back to Wake over a 70-sec period in (**A**) male and (**B**) female *orexin/tTA; TetO-DTA* mice during Week 6 in the DOX(-) condition. Example EEG spectrograms (7 10-sec epochs), EEG and EMG traces illustrating typical transitions from Wake to the Delta State back to Wake over a 70-sec period in (**C**) male and (**D**) female *orexin/tTA; TetO-DTA* mice during Week 6 in the DOX(-) condition. Raw EEG power during the first 6-h of the dark period for the delta (0.5-4 Hz), theta (6-9 Hz), alpha (9-12 Hz), beta (12-30 Hz), low gamma (30-60 Hz) and high gamma (60-100 Hz) bandwidths during Cataplexy, the Delta State, NREM, REM and Wake during Week 6 in the DOX(-) condition for a male (**E**) and a female (**F**) *orexin/tTA; TetO-DTA* mouse. Insets show raw power in the 0-10Hz range, plotted as a rolling average of three adjacent 0.122 Hz bins. (**G)** Total time in the Delta State during the first 6-h of the dark period during baseline (Week 0) and Weeks 1, 2, 4 and 6 in the DOX(-) condition during which time the Hcrt neurons are expected to degenerate. (**H)** Mean Delta State bout duration and (**I**) number of Delta State bouts during the first 6-h of the dark period during baseline (Week 0) and Weeks 1, 2, 4 and 6 in the DOX(-) condition. Values in panels E – I are mean ± SEM. * above the bars in panels E-I indicate significance (*p* < 0.05) as determined by Dunnett’s *post hoc* test. For E and F, spectral power for DS within each bandwidth was compared to all other states by 1-way ANOVA and mixed-effects analysis.

Both male and female mice exhibited DS which progressively increased as the degeneration progressed, primarily due to an increased number of DS bouts (**Figures 9G-9I**). EEG power spectra during DS were similar to the NREM power spectra (**Figures 9E, 9F**). In both sexes, delta power was greater in DS compared to W, REM or C, spectral power in the theta, alpha and beta bands was greater in DS than in W, and beta power was also greater in DS than in cataplexy and REM sleep. Theta power in DS was lower than during cataplexy in both sexes. No significant differences in any of the frequency bands were found between DS and NREM. Therefore, the EEG power spectra of DS was clearly different from that found during REM, cataplexy or Wake and most resembled that of NREM sleep.

## Discussion

Although the consequences of dietary DOX withdrawal and subsequent Hcrt neuron degeneration has been well-documented in male *orexin-tTA; TetO DTA* mice^30^, similar information for female mice of this strain has been lacking. This is not unusual as, with some notable exceptions^35-42^, female mice have been rarely used in sleep studies^27,43^. The primary symptoms of human narcolepsy that can be assessed in a mouse model of this disorder are excessive daytime sleepiness (EDS), nocturnal sleep disruption and cataplexy, the pathognomonic symptom of NT1.

We find that female DTA mice exhibit cataplexy sooner after DOX removal than their male counterparts: by Week 1, all female mice exhibited cataplexy whereas only 3 of 7 male mice showed this behavior. In contrast, all mice of both sexes exhibited cataplexy by Week 2 and cataplexy levels were similar in both sexes by Week 6. As described for males^30^, cataplexy in females primarily occurred in the dark (active) phase, although some cataplexy also occurs during the light (inactive) phase in both sexes (**Figure 2O vs. 2P**). In males, mean cataplexy bout duration during the dark phase increased with time post-DOX removal as noted previously^30,41^, but this relationship was not evident for female DTA mice. The amount of cataplexy during the 12-h dark phase has previously been reported to be similar in males and females^41^, results that are congruent with our findings in Weeks 2-6. However, the same study reported that female DTA mice exhibit more cataplexy bouts in the 12-h dark period than males^41^, which differs from our results (**Figure 5G**). Inter-laboratory differences in substrain, housing conditions (presence or absence of a running wheel) or scoring of cataplexy may have a role here as the mean number of cataplexy bouts during the dark phase (**Figure 5G)** is more than twice the number reported previously for female DTA mice^41^. The more rapid emergence of cataplexy in females along with the absence of the progressive increase of cataplexy duration observed in males suggests that Hcrt neurons may degenerate faster in female than in male DTA mice. Alternatively, the neural circuits dependent upon input from Hcrt neurons may be more sensitive in females than males.

The progressive decrease in Wake Bout Duration and increased number of wake bouts in both male and female DTA mice are indicative of disrupted sleep/wake architecture as Hcrt neuron degeneration proceeds. Both sexes exhibited 6-7 fold more wake bouts during the dark in Week 6 compared to baseline, indicating an inability to sustain wakefulness that was also reflected in the progressive reduction in mean wake bout duration. Together, these two parameters mirror EDS in people with narcolepsy for whom daytime naps are not only common, but a necessity for some individuals. It is interesting to note here that, although cataplexy bout duration in female DTA mice reaches a plateau by Week 1, wake bout duration progressively decreases with weeks after DOX removal. These results indicate that, in females, a more limited disruption of Hcrt neurotransmission results in cataplexy whereas the severity of EDS symptomatology may be more dependent on the extent of Hcrt neurotransmission disruption.

The present study provides limited evidence that the nocturnal sleep disruption characteristic of people with narcolepsy is reflected in DTA mice. In male DTA mice, there was a reduction of mean NREM bout duration during the light phase in Week 6 (**Figure 4D)** and there is a comparable non-significant trend in females, but the number of NREM bouts is constant in both sexes as Hcrt neuron degeneration proceeds (**Figure 5D**). Transient increases in mean REM bout duration occur during the light phase in both sexes as Hcrt neuron degeneration proceeds (**Figure 4F**) but these changes are not sustained.

As mentioned above and as illustrated by the large variation in wake bout duration in females (**Figure 4A**), one female DTA mouse remained awake throughout the entire dark (active) phase during baseline Week 0. Female C57BL/6 mice are known to spend less time awake during the dark period than males^35,39^ but, even in comparison to the female mice reported in those studies, this DTA female mouse is an outlier. Although the estrus cycle is reported to have minimal effects on sleep parameters relative to background strain^42^, to control for any effect of the estrus cycle on sleep/wake, we conducted the video/EEG/EMG recordings at 4 day intervals. The sustained wakefulness of this female at baseline contrasts with the more typical pattern observed from this same individual during the dark phase in the Week 1, 2, 4 and 6 recordings during which this female should have been at the same stage of the estrus cycle. Moreover, none of the other mice recorded simultaneously with this mouse during Week 0 showed excessive wakefulness during the dark phase.

### Reduced T_b_ rhythm as Hcrt degeneration proceeds

The T_b_ rhythm reported by the subcutaneously-placed DSI transmitters clearly shows a decline in the T_b_ rhythm as Hcrt degeneration proceeds. Although this reduction is most evident in males (**Figure 6C, E**), ANOVA indicated that this decline was significant compared to the baseline (Hcrt neuron intact) condition in both sexes from Weeks 2-6. A study of human narcoleptics (conducted prior to the distinction between NT1 and NT2) that selected experimental subjects on the basis of short REM sleep onset found that rectal T_b_ was higher in people with narcolepsy at night than in control subjects, which was attributed to the disturbed nocturnal sleep of the patients.^44^ Similarly, *prepro-orexin* knockout (KO) mice have a higher core T_b_ during sleep than wild type (WT) mice, which was attributed to either a deficiency in the heat loss or sustained activity of heat production mechanisms.^45^ On the other hand, Hcrt/orexin neuron-ablated mice failed to exhibit the increased T_b_ that occurs in WT and KO mice in response to handling stress^46^, implicating a colocalized Hcrt neuron neurotransmitter in thermoregulatory responses. Taken together, these data suggest a role for Hcrt neurons, although not the peptides themselves, in thermoregulation. Since mice of different strains and sexes prefer temperatures between 26-29°C^47^ but our experiments were conducted at an ambient temperature of 22 ± 2ºC, the DTA mice may have been unable to maintain T_b_ at the baseline level while Hcrt neuron degeneration was proceeding. Human narcoleptics show an abnormal distal to proximal skin temperature gradient while both awake and asleep, suggesting a role for Hcrt/orexin neurons in skin temperature regulation^48^. Furthermore, manipulation of distal to proximal skin temperature improved nocturnal sleep in people with narcolepsy^49^.

### EEG Spectra and Delta State

The EEG spectra for waking and cataplexy reported in **Figures 7 and 8** are based on human-scored EEG/EMG/video recordings collected from the first 6-h of the dark period (ZT13-18) and excluded epochs that contain artifacts that could otherwise contaminate the analyses. Consistent with the concept of narcolepsy as an arousal state boundary disorder, the waking EEG spectra in both sexes of DTA mice are characterized by progressively elevated delta, theta and beta frequencies as the Hcrt neurons degenerate (**Figure 7**). Whereas some of these increases may be transient, the increased beta activity in female DTA mice persists at least as long as 6 weeks post-DOX. The waking EEG spectra show further evidence of a slower progression of the development of the narcoleptic phenotype in males compared to females. The normalized wake power between 0.5-40 Hz for Week 1 is distinctly different from Weeks 2-6 in males but not in females (**Figures 7A, 7B**).

For the EEG spectra during cataplexy, statistical analyses compared to the DOX(+) condition could not be performed since no cataplexy occurred on Week 0; consequently, our interpretation of the EEG spectra during cataplexy is qualitative (**Figure 8**). Overall, the EEG spectra in cataplexy was very similar between males and females and also showed a similar progression, with lower levels of theta (**Figure 8D**) and beta (**Figure 8F**) power in Week 1 than in subsequent weeks.

Careful analysis of the EEG, EMG and video recordings by our expert scorers revealed a unique state that we called the DS, which was observed in both sexes but only while the Hcrt neurons were degenerating. DS events were typically ∼30 sec in duration, ranging from 1 to 6 10-sec epochs, and were both preceded and followed by active Wake. Although the mice ceased movement during DS as in cataplexy, muscle tone was maintained but phasic EMG activity was absent. The EEG was synchronized with high delta activity as in NREM sleep but, as evident from the video recordings which were essential to recognize DS, the mice remain standing with their eyes open. The delta power during DS in both sexes was greater than that observed in Wake, REM and Cataplexy and the theta power was less than that observed during REM sleep (**Figure 9E, 9F**). DS was distinguished from Wake not only by the increased delta power but spectral power in the theta, alpha and beta bands was greater in both sexes during DS compared to Wake. DS has some similarities to “delta-theta sleep” which we recently described in mice in which both the Hcrt and melanin-concentrating hormone (MCH) neurons were ablated.^50^ While we find increased delta in DS, we did not find increased theta as has been reported for delta-theta sleep^50^. However, methodological differences, such as different electrode configurations, could be a confound to preclude a direct comparison between DS and delta-theta sleep. Unfortunately, we did not follow this cohort of mice beyond 6 weeks DOX(-), so we do not know whether DS is a transitional state that only occurs while the Hcrt neurons are degenerating or whether this state occurs throughout the remainder of the mouse’s life. Although the DS state bears some resemblance to both “sleep attacks” and microsleeps, the exact analog of DS in humans remains to be unequivocally identified. Nonetheless, DS appears to be one more manifestation of the arousal state boundary instability that characterizes narcolepsy in both humans and animal models of this disorder.

### Use of Somnivore

In this study, we utilized the machine learning-based (ML) program Somnivore^34^ to aid in scoring the remaining 18-h of the 24-h recordings for both male and female mice. Somnivore has been validated in a variety of eutherian mammals including birds^51^, mice^34,52^, rats^34^ and humans^34^ and has also been used in another mouse model of narcolepsy, the *prepro-orexin* KO mouse^34^. The use of automated scoring systems for classification of arousal states is becoming increasingly widespread and has recently been implemented in studies of DTA mice^41^. Once a user has been trained in Somnivore, a 24-h EEG/EMG recording of a DTA mouse can be scored in 20-30 min. Thus, this tool and others like it hold the promise of greatly expediting sleep studies in rodents, in particular, although trained users must always be vigilant to monitor the raw recordings for unusual states such as DS.

### Limitations of the present study

Although our study is a thorough characterization of male vs. female DTA mice, it is not without its limitations. First, our conclusions are based on a relatively small N for each sex which limits some statistical analyses yet, by and large, the results appear robust. As indicated above, we have utilized a ML-based algorithm to score the last 18-h of 24-h recordings, including cataplexy. Since video-based information is yet to be incorporated into this algorithm, the identification of cataplexy during the last 18-h was based on a training set of epochs scored as cataplexy by expert scorers who utilized the videos collected during the first 6-h of the 24-h recording. Nonetheless, all epochs scored as cataplexy by Somnivore were re-inspected by expert scorers who had access to the video recordings for epochs classified by Somnivore as cataplexy. Although this quality control step was absolutely crucial to the entire process, it was somewhat laborious. Despite such efforts, our study did not replicate the results of a recent publication^41^ which claimed that REM sleep was elevated in male DTA mice during the dark phase. Although there are many procedural differences between these two studies that could account for this difference, one possible source that cannot be ignored is our use of a ML algorithm to score the second half of the dark phase.

### Conclusions and Perspective

Although female DTA mice develop cataplexy, fragmented wakefulness, and EEG spectral changes more rapidly than their male counterparts, there appear to very few differences between the two sexes by 6 weeks DOX(-). Accordingly, we advocate for use of both sexes of this unique mouse model of narcolepsy to ensure that information relevant to the entire human population is acquired and that the goal of reducing the use of the number of animals can be achieved. Furthermore, to expedite sleep studies, we find that a ML-based algorithm can be utilized with high confidence by expert scorers as long as they remain vigilant for atypical EEG states. Lastly, we have identified a unique state called DS that resembles sleep attacks that occur in human narcolepsy.

As suggested above, the earlier occurrence of cataplexy as well as the absence of the progressive increase of cataplexy duration in females suggests a topic for future research. Do the Hcrt neurons degenerate faster in female than in male DTA mice and/or are the neural circuits in females more sensitive to loss of Hcrt input than in males? Further definition of the DS, including its similarities and differences with delta-theta sleep, as well as identification of the neural substrates that underlie this state is likely to be a productive future research area.

## Acknowledgements

Research supported by NIH R01 NS098813 and R01 NS103529 to T.S.K. We thank Haley Courtney, Laure Alexandre and Sahin Ozsoy for technical assistance and Chihung (Jeffrey) Hung for his comments on the distinction between the Delta State and Delta-Theta Sleep described in ^50^.

